# IL-7 is essential for accumulation of antigen-specific CD8 T cells and to generate clonotype-specific effector responses during airway influenza/A infection

**DOI:** 10.1101/2021.04.21.440748

**Authors:** Abdalla Sheikh, Jennie Jackson, Hanjoo Brian Shim, Clement Yau, Jung Hee Seo, Ninan Abraham

**Affiliations:** Department of Microbiology and Immunology, University of British Columbia, Vancouver, Canada; Life Sciences Institute, University of British Columbia, Vancouver, Canada; Calvin, Phoebe and Joan Snyder Institute for Chronic Diseases, University of Calgary, Alberta, Canada; Duke-NUS Medical School, 8 College Road, Singapore; Department of Zoology, University of British Columbia, Vancouver, Canada

**Author notes:** Corresponding author: Ninan Abraham.

## Abstract

Airborne diseases are the leading cause of infectious disease-related deaths in the world. In particular, the influenza virus activates a network of immune cells that leads to clearance or an overzealous response that can be fatal. Tight regulation of the cytokines that enable proper activation and function of immune cells is necessary to clear infections efficiently while minimizing damage to the host. Interleukin-7 (IL-7) is a cytokine known for its importance in T cell development and survival. How IL-7 shapes CD8 T cell responses during an acute viral infection is less understood. We had previously shown that IL-7 signaling deficient mice have reduced accumulation of influenza-specific CD8 T cells following infection. We sought to determine whether IL-7 affects early CD8 T cell expansion in the mediastinal lymph node and effector function in the lungs. Using IL-7Rα signaling deficient mice, we show that IL-7 is required for a normal sized mediastinal lymph node and the early clonal expansion of antigen-specific CD8 T cells therein. Bone marrow chimeric models and adoptive transfer of transgenic TCR CD8 T cells reveal a cell-intrinsic role for IL-7 in the accumulation of NP_366–374_ and PA_224–233_-specific CD8 T cells. We also found that IL-7 dictates terminal differentiation, degranulation and cytokine production in PA_224–233_-specific but not NP_366–374_-specific CD8 T cells. We further demonstrate that IL-7 is induced in the lung tissue by viral infection and we characterize multiple cellular sources that contribute to IL-7 production. Drugs that manipulate IL-7 signaling are currently under clinical trial for multiple conditions. Our findings on IL-7 and its effects on lower respiratory diseases will be important for expanding the utility of these therapeutics.

**Author Summary:** Interleukin-7 plays an important role in development of immune cells such as lymphocytes. In recent years, its role in the immune system has been expanded beyond the development of immune cells to include revitalizing of lymphocytes during tumor and chronic viral response. We show here that IL-7 is required for accumulation and function of specialized lymphocytes in the lungs during an acute influenza infection.

## Introduction

The influenza virus is an airway pathogen that infects lung epithelial cells and activates a network of immune cells. It causes seasonal and pandemic outbreaks with major global health and economic impacts. Seasonal variants of influenza can cause death in children, the elderly and immune-compromised individuals (1). Vaccination is a cornerstone of the preventative measures taken towards influenza as it arms the adaptive immune system. Multiple cell types in the immune system are required during a response to influenza infection. However, we lack a complete understanding of the cellular aspects and intercellular signaling components that lead to efficient generation of functionally competent immune cells. At the center of immune responses are the cytokine signals that shape various aspects of immune cells (2).

A hallmark of our immune response is its ability to develop memory to previously encountered pathogens – T cells are major players in this process. An ideal anti-viral response to influenza and other viruses, requires cytotoxic CD8 T cells for their swift and specific response. CD8 T cells employ multiple methods to kill infected cells and control viral replication, namely, granule-dependent (granzyme B and perforin) and ligand-dependent (Fas-FasL) means (3).

In addition to TCR-MHC engagement (signal 1) and co-stimulation (signal 2), cytokine cues (signal 3) have great influence in activating and shaping CD8 T cell responses and their terminal differentiation. Once a CD8 T cell receives these signals, it is driven towards a robust clonal expansion phase whereby a single cell expands to ∼10^5^ cells (4). The signal 3 cytokines that govern T cells are multifaceted and include interleukin-2 (IL-2), IL-6, IL-10, IL-12, IL-15 and others which dictate their terminal differentiation and inflammatory functions (5, 6).

The common gamma chain (γc) cytokine IL-7 is produced mainly by stromal cells in the bone marrow and thymus. At steady state, it plays an indispensable role in the development of both pre- and pro-B cells and T cells (7-10). IL-7 is important in the development and survival of T cells at specific stages of maturation in the thymus as the expression of IL-7Rα (CD127) is dynamically regulated (8, 11-13). IL-7 shares the IL-7Rα with thymic stromal lymphopoietin (TSLP), an alarmin cytokine that plays a major role in mucosal sites. In addition to its role in development, IL-7 also plays a canonical role in maintenance of memory T cells (14). The span of IL-7’s function was further expanded in the past decade when it was shown to be able to shape the effector responses of cytotoxic CD8 T cells by enhancing their responses against tumors (15) and bacterial infection (14), and reverse T cell exhaustion caused by chronic LCMV infection, thus, preventing liver pathology (16). However, the extent to which IL-7 regulates CD8 T cell response to acute viral infections is unknown. We had previously shown that IL-7 but not TSLP is important for the accumulation of influenza-specific CD8 T cells in the lungs but the mechanism by which this occurs is unclear (17). Since IL-7 is implicated in over 8 clinical trials for treatment of infections, solid tumors and other chronic conditions, the intricacies of IL-7 signaling in functional outcomes requires further inquiry (18).

In this study, we asked: what modulatory effects does IL-7 have on CD8 T cell priming and effector functions during an acute airway influenza infection? Using IL-7Rα knock-in mice, we have shown that in the lung draining mediastinal LNs (mdLNs), IL-7 is important for early priming and accumulation of CD8 T cells specific for influenza NP_366-374_ and PA_224-233_ presented on H2D^b^ in a cell intrinsic manner. We also show that IL-7 is important for the terminal differentiation and cytokine production in CD8 T cells. This study will aid in therapeutic development and vaccine adjuvant studies to design combinatorial therapeutic strategies.

## Results

### IL-7Rα signaling is required for accumulation of influenza-specific CD8 T cells

To assess the importance of IL-7 signaling in CD8 T cells following infection with influenza, we infected WT and IL-7Rα^449F^ mice with A/PR/8/34 (PR8) influenza virus and measured influenza-specific CD8 T cells by flow cytometry using MHC-I tetramers. We found that IL-7Rα^449F^ mice have reduced proportions of NP_366-374_ and PA_224-233_-specific cells within CD8 T cells in the lungs 7 days post-infection (dpi) (Fig. 1a), which phenocopies past observation of this defect at 9 dpi in a new, embryo re-derivation based, specific pathogen free facility (17). Since the majority of pathogen-specific T cells originate from tissue draining lymph nodes, we examined the mediastinal lymph nodes (mdLNs) of infected mice and found that IL-7Rα^449F^ mice have reduced lymph node sizes, particularly the mdLN, compared to WT mice. Interestingly, unlike WT mice, there was little increase in mdLN size of IL-7Rα^449F^ mice after influenza infection (Fig. 1b). Additionally, enumeration of the total antigen-specific cells in the mdLN at multiple days post infection revealed a consistent and substantial defect that is not due to a delay in expansion kinetics. (Fig. 1c). Consistent with lack of LN hyperplasia, the proportion of antigen-specific cells in IL-7Rα^449F^ mdLN at 5 dpi was reduced indicating a defect in early priming of CD8 T cells (Fig. 1d).

**Figure 1.**
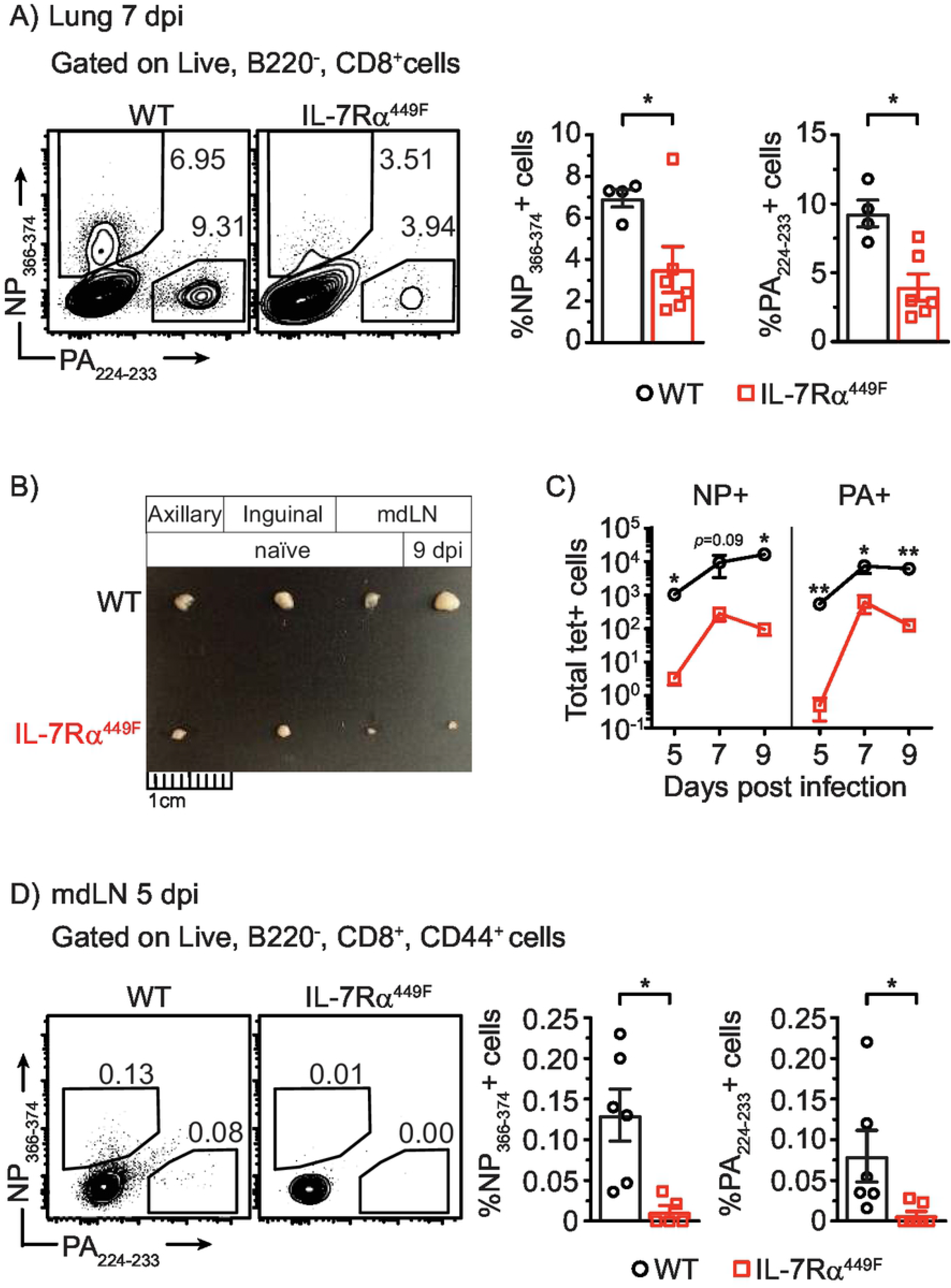
Accumulation of tetramer specific response in IL-7Rα^449F^ is impaired following influenza infection. **(a, b)** Representative FACS plots and bar graphs of the frequency of NP_366-374_^+^ and PA_224-233_^+^ cells within CD8 T cells in **(a)** the lungs 7 dpi and **(b)** mdLN 5 dpi of WT and IL-7Rα^449F^ mice. Gated within Live B220^-^, CD8^+^, (CD44^+^) cells. **(c)** Photograph images offering comparison of various mouse lymph nodes and **(d)** total number of NP_366-374_^+^ and PA_224-233_^+^ cells in the mdLN of WT and IL-7Rα^449F^ mice at the indicated days post infection. Data are representative of 2–3 experiments with n=4–7 per genotype. \**P*<0.05 as determined by two-tailed Student’s t-test.

### Intrinsic requirement for IL-7Rα signaling in the accumulation of influenza-specific CD8 T cells in the mdLN

Previous reports have shown that IL-7 is required for the generation of lymph nodes independent of the peripheral lymphocyte pool (19, 20). This likely contributed to the reduced lymph node sizes noted above. Therefore, it is possible that the reduction in influenza-specific CD8 T cell accumulation was due to factors extrinsic to T cells in the lymph node and that IL-7 was indirectly important for shaping the cellular and cytokine environment for optimal T cell activation. To address this, we created bone marrow (BM) chimeric mice whereby we grafted BM cells of wild type (WT) and IL-7Rα^449F^ mice into lethally irradiated RAG-1-deficient hosts (Fig. 2a). Since WT lymphocytes outcompete IL-7Rα^449F^ lymphocytes during development (21), we delivered a 1 to 10 ratio of WT to IL-7Rα^449F^ cells respectively. Following engraftment and infection of the hosts, we noted a reversal of this ratio within the CD8 T cell compartment in the mdLN (Fig. 2b). More importantly, IL-7Rα^449F^ CD8 T cells resulted in reduced NP_366-374_ and PA_224-233_ - specific cells in proportion despite engraftment in a competent niche (Fig. 2b). These data suggest that IL-7Rα signaling plays an intrinsic role necessary for CD8 T cell expansion during influenza infection.

**Figure 2.**
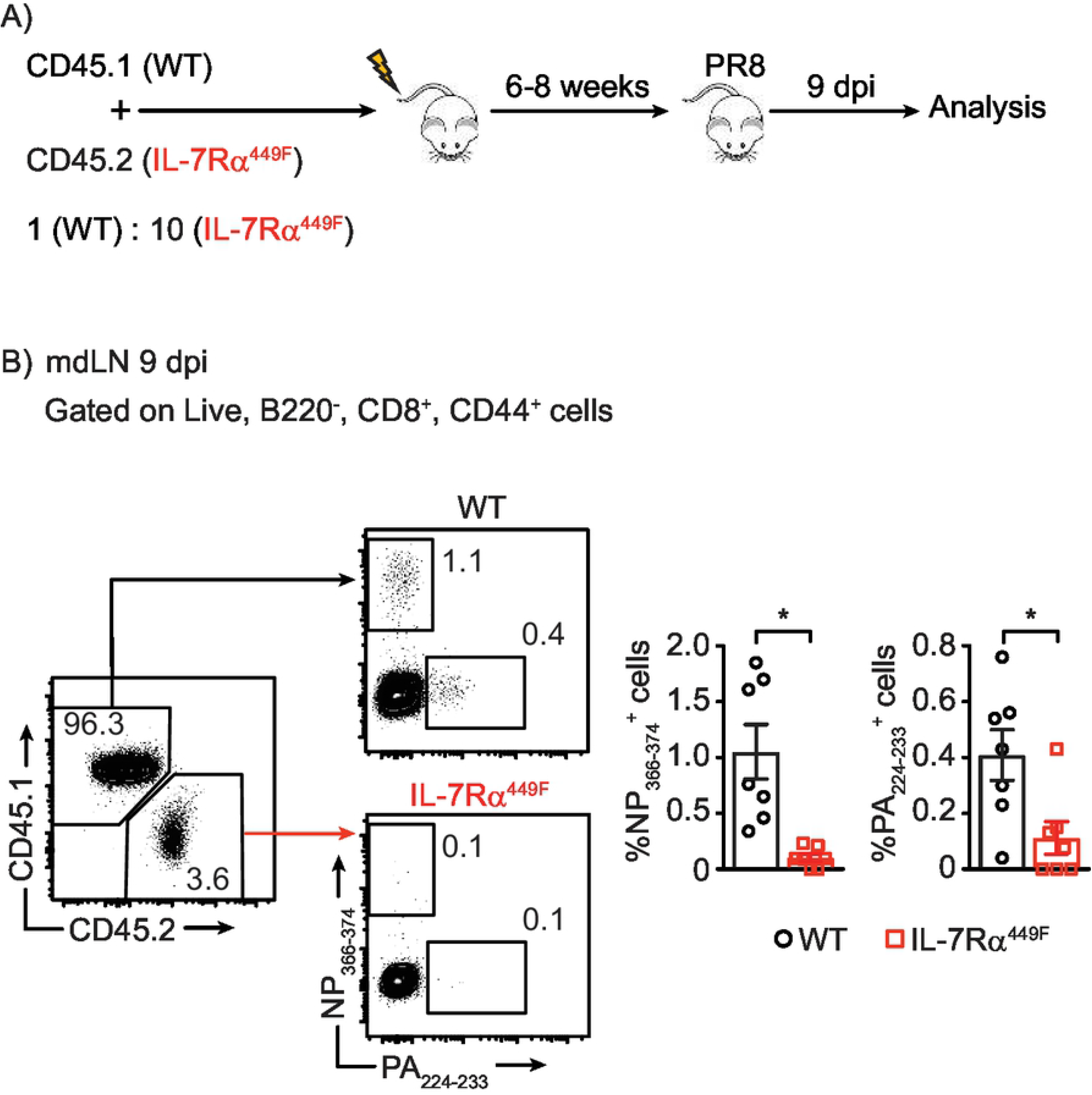
Impairment in tetramer specific response seen in the mdLN of IL-7Rα^449F^ mice is cell intrinsic. **(a)** Schematic of bone marrow chimera set-up. **(b)** Representative FACS plots and bar graphs of the frequency of NP_366-374_^+^ and PA_224-233_^+^ cells within CD8 T cells in the mdLN of WT and IL-7Rα^449F^ chimeric mice 9 dpi. Gated within live, B220^-^, CD8^+^, CD44^+^, CD45.1^+^ or CD45.2^+^ cells. Data are representative of two experiments with n=5-7 per genotype. \**P*<0.05 as determined by two-tailed Student’s t-test.

### IL-7Rα plays a role in early priming of CD8 T cells independent of TCR and number of naïve precursors in the mdLN

To determine if the reduction in pathogen-specific CD8 T cells was due to reduced numbers of naïve precursors or a result of gaps in TCR repertoire, we adoptively transferred SIINFEKL OVA peptide-specific and MHCI restricted transgenic TCR CD8 T cells from OT-I mice crossed with IL-7Rα^449F^ mice (CD45.2) in to BoyJ mice (CD45.1). We infected these mice with a modified version of the influenza PR8 virus that has the OVA (SIINFEKL) peptide inserted into the stalk of the NP polypeptide (influenza PR8-OVA). Despite delivering equal number (1×10^6^) of OT-I and OT-I;IL-7Rα^449F^ CD8 T cells into distinct BoyJ (CD45.1) hosts, OT-I;IL-7Rα^449F^ CD8 T cells did not expand to the same extent as wild type OT-I CD8 T cells 4 dpi (Fig. 3a) in the mdLN. This defect was observed as early as 3 dpi (Suppl. Fig. 1). Interestingly, the expression of the early activation marker CD5 at 4 dpi was significantly reduced in OT-I;IL-7Rα^449F^ CD8 T cells indicating a defect in priming (Fig. 3b and c). Furthermore, TCR expression on OT-I;IL-7Rα^449F^ CD8 T cells showed a trend towards higher expression at 4 dpi albeit not significantly, further suggesting a possible defect in early priming (Fig. 3b and c).

**Figure 3.**
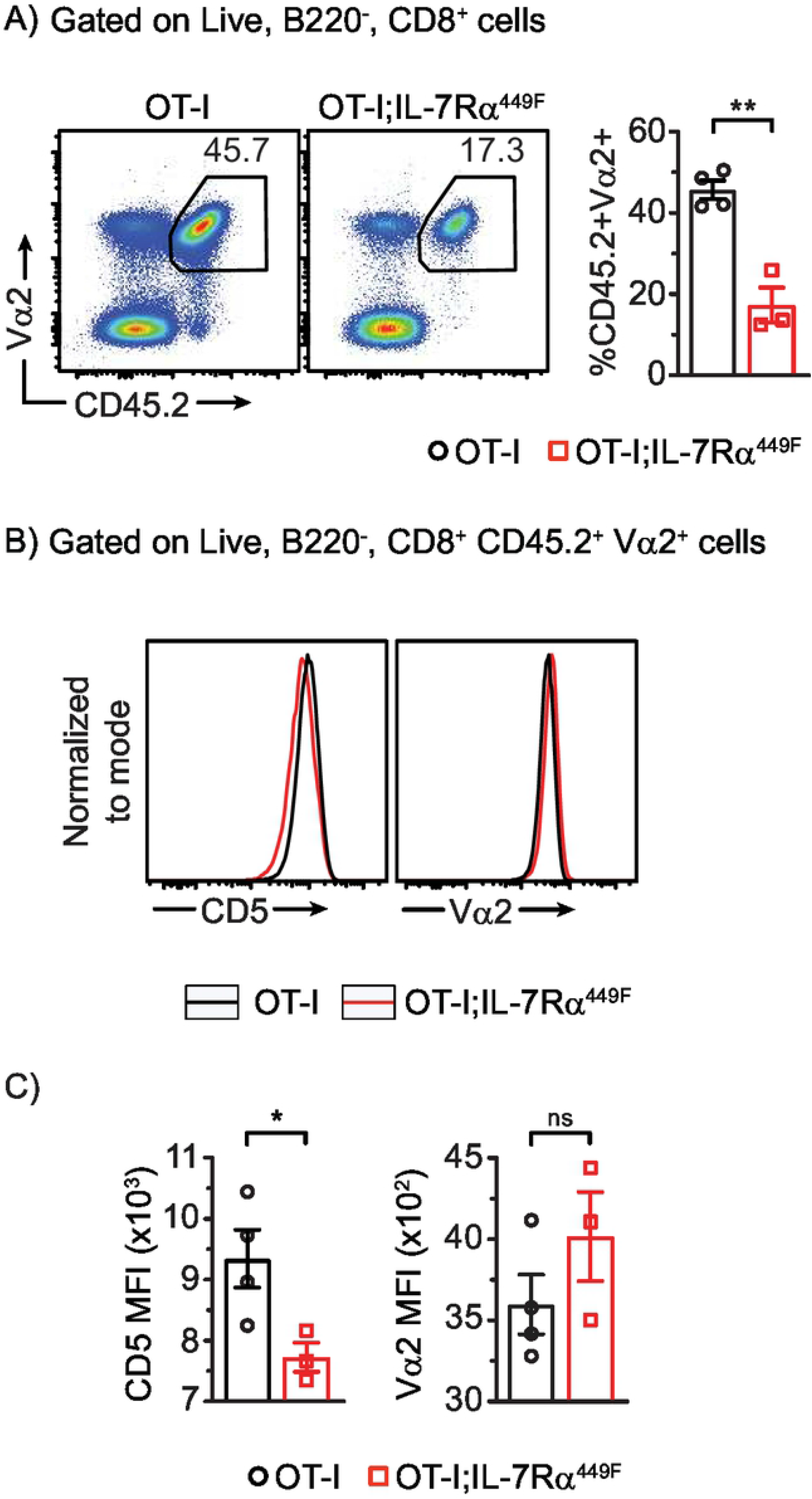
Expansion of adoptively transferred OTI-IL-7Rα^449F^ CD8 T cells is impaired in the mdLN following influenza infection. **(a)** Scatter plot and representative bar graph of CD45.2^+^ Vα2^+^ CD8 T cells in BoyJ (CD45.1^+^) mice 4 dpi. Gated within live B220^-^ CD8^+^ cells. **(b)** Histogram and **(c)** bar graph of median florescence intensity (MFI) of activation markers (CD5, TCR (Vα2), and CD69). Data are representative of two experiments. \**P*<0.05 and \*\**P*<0.01 as determined by two-tailed Student’s t-test.

### Increased dendritic cell accumulation in the lungs of IL-7Rα^449F^ mice

Evidence of IL-7 cell-intrinsic effects on CD8 T cells does not exclude cell-extrinsic effects. Antigen presenting cells, specifically dendritic cells (DCs) are key to activation of CD8 T cells and their subsequent response. Previous reports have demonstrated that IL-7Rα signaling plays an indirect role in the development of conventional DCs (22). Furthermore, IL-7 has been shown to regulate CD4 T cell proliferation in conditions of lymphopenia indirectly though DCs (23). We found that in the lungs of IL-7Rα^449F^ mice, CD11b^+^ CD103^-^ DCs but not CD11b^-^ CD103^+^ DCs, accumulate up to 9 dpi, while in WT mice these DCs peak at 7 dpi and decrease in numbers at 9 dpi (Suppl. Fig. 2a). This increased accumulation could be due to a cell-extrinsic factors such as increased viral load in the IL-7Rα^449F^ mice as a result of lack of appropriate T cell response. Another plausible reason could be due to impaired lymph node homing signals from chemokines as a result of reduced draining lymph node size. Alternatively, IL-7 may have a direct effect on CD11b^+^ CD103^-^ DC maturation or migration. To test these hypotheses, we created 50:50 BM chimeras using WT:IL-7Rα^449F^ or WT:WT BM cells (CD45.1:CD45.2) grafted into congenic BoyJ/WT hosts (CD45.1/.2). We found that after infection, WT:IL-7Rα^449F^ ratios of DC subsets were comparable to WT:WT ratios in both the mdLN and lungs for both DC subsets (Suppl. Fig. 2b and c). This suggests that the phenotypic elevation of DCs in IL-7Rα^449F^ mice during influenza infection has a cell extrinsic cause.

### IL-7 is inducible in lung tissues in response to influenza

IL-7 is mainly produced by radio-resistant cells such as stromal and epithelial cells of the bone marrow and thymus, where it plays a major role in hematopoiesis and thymopoiesis (10, 24). A few studies have demonstrated IL-7 expression in various tissues including liver, skin, intestines and lungs (25-29). While IL-7 is mainly produced in steady state lungs by lymphatic endothelial cells, its source during inflammation is unclear (30, 31). To assess IL-7 expression dynamics in response to influenza, we first infected human type II epithelial cells (A549) with influenza and assessed *Il7* mRNA using qRT-PCR. We found that IL-7 expression is induced within 24h following influenza infection correlating with the antiviral response signified by IFN-β and viral replication demonstrated by M1 mRNA transcript expression (Fig. 4a).

**Figure 4.**
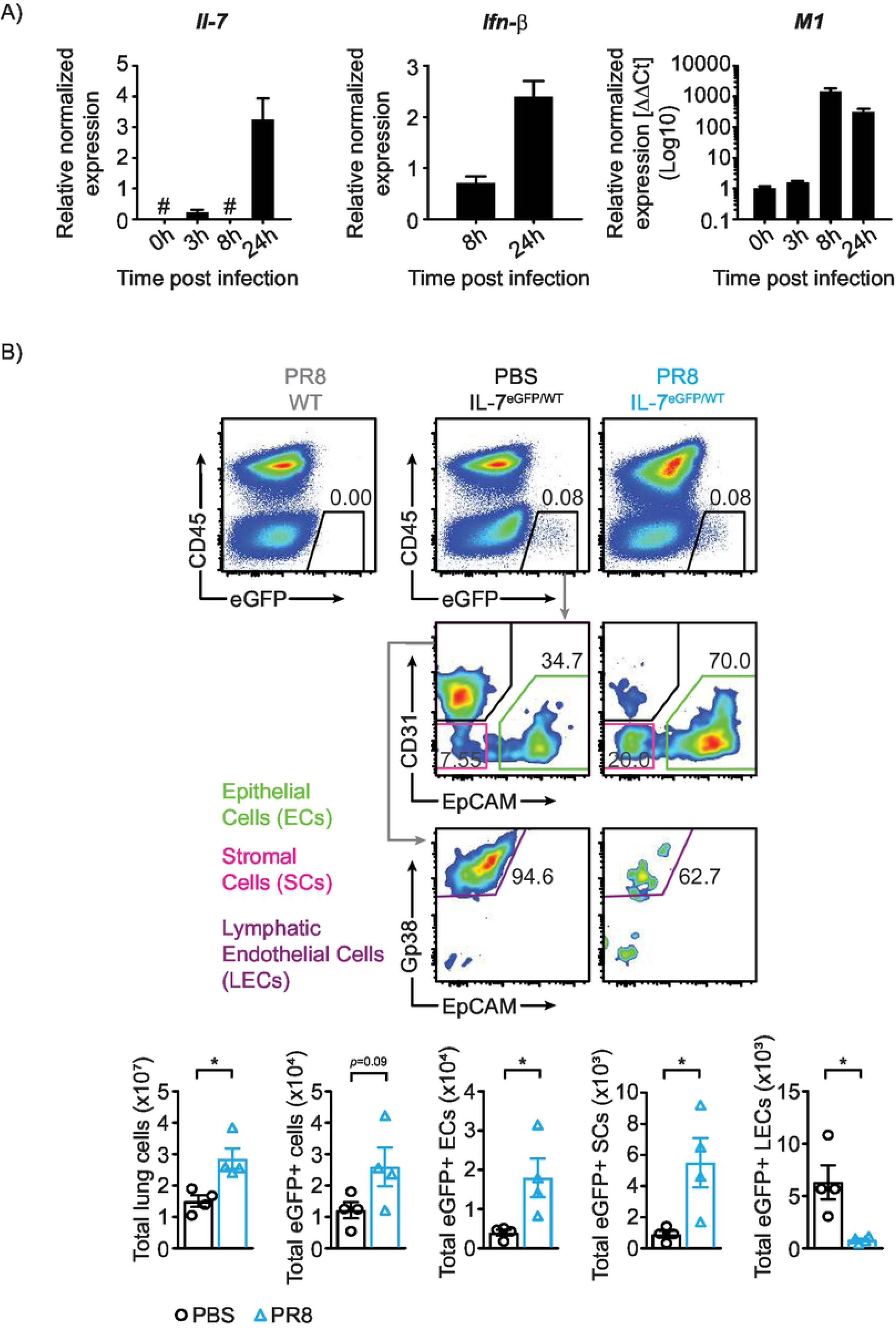
IL-7 expression in lung tissues. **(a)** Quantitative PCR of IL-7, IFN-β and M1 in A549 cells at the indicated times post infection normalized against beta actin. M1 expression is further normalized to time point 0h. Data is representative of 2 experiments n=3 per experiment. **(b)** Expression of IL-7 in various CD45^-^ lung cells using IL-7^eGFP/WT^ mice. Epithelial cells (ECs) are CD45^-^ EpCAM^+^, stromal cells (SCs) are CD45^-^ EpCAM^-^ CD31^-^ and lymphatic endothelial cells (LECs) are CD45^-^ EpCAM^-^ and CD31^+^ GP38^+^. Data are representative of two experiments with n=4 per genotype. \**P*<0.05 as determined by two-tailed Student’s t-test.

To determine if IL-7 is induced *in vivo*, we used the IL-7^eGFP/WT^ mice. We noted an increase in the number of cells expressing eGFP at 6 dpi (Fig. 4b). Interestingly, the majority of IL-7-eGFP^+^ cells during infection are epithelial cells. The number of IL-7-eGFP^+^ stromal cells also significantly increased during infection. However, IL-7-eGFP^+^ lymphatic endothelial cells were reduced following infection (Fig. 4b). Altogether, these *in vitro* and *in vivo* experiments suggest that lung epithelial cells are responsive to viral infection, and that during influenza infection, they become the primary source of IL-7.

### IL-7Rα^449F^ CD8 T cells have reduced terminal differentiation

The cytokine milieu that CD8 T cells are exposed to throughout the course of an immune response governs their terminal differentiation to effector cells and hence, their functional capabilities. Among the heterogeneous population of CD8 T cells that emerge during the expansion phase are the T-bet^hi^ and granzyme B producing short-lived effector cells (SLECs) that are identified by their expression of the killer cell lectin-like receptor G1 (KLRG1) and low CD127 (IL-7Rα) (6, 32). Due to extra-physiological expression of the mutated IL-7Rα^449F^ subunit (unpublished data), we limited our use of SLEC markers to KLRG1. Testing the expression of KLRG1 in peptide-specific cells revealed a reduced proportion of KLRG1^+^ cells in both NP_366-374_ and PA_224-233_ - specific cells of IL-7Rα^449F^ mice (Fig. 5). However, the difference in KLRG1 expression between WT and IL-7Rα^449F^ mice was greater in PA_224-233_-specific compared to NP_366-374_-specific cells (Fig. 5). These data suggest that IL-7Rα signaling plays a role in the terminal differentiation of antigen specific CD8 T cells.

**Figure 5.**
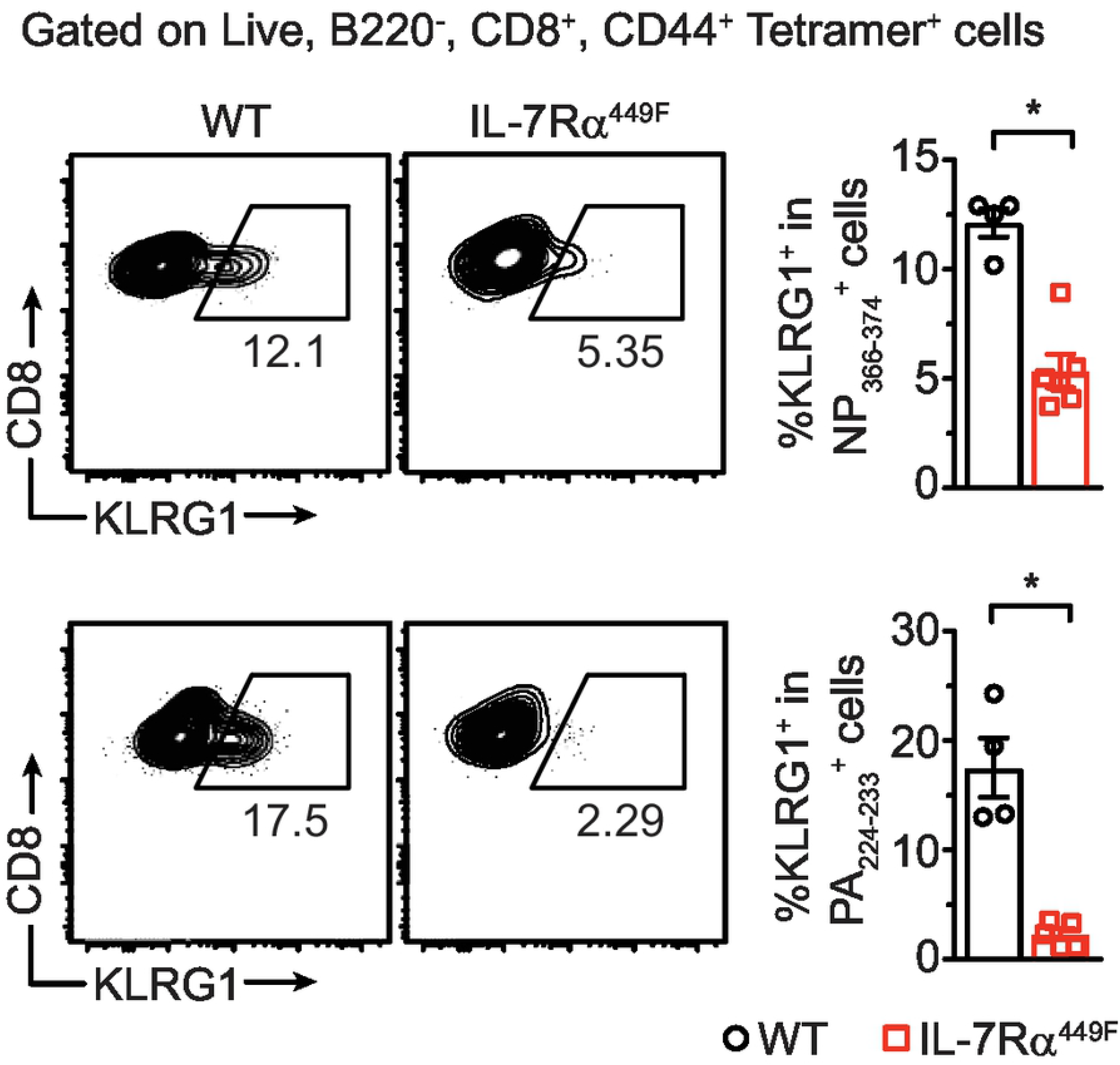
CD8 T cells of IL-7Rα^449F^ mice have reduced terminal differentiation. Scatter plots and bar graphs showing flow cytometric analysis of KLRG1 expression as a percentage within lung NP_366-374_+ and PA_224-233_+ CD8 T cells of WT and IL-7Rα^449F^ mice 7-9 dpi. Data are representative of three experiments with n=4–6 per genotype. \**P*<0.05 as determined by two-tailed Student’s t-test.

### Reduced degranulation and cytokine production by IL-7Rα^449F^ CD8 T cells

Secretion of cytotoxic granules and inflammatory cytokines is a major event in the CD8 T cell effector response. Lysosome associated membrane protein-1 (LAMP-1) or CD107a is a membrane glycoprotein found in the lumen of granzyme B and perforin containing vesicles. Detection of CD107a on the surface of CD8 T cells through flow cytometry provides a direct method for identifying degranulating cells (33). Using this method, we noted that CD8 T cells of IL-7Rα^449F^ mice had reduced CD107a expression in antigen experienced cells (CD44^+^) and in PA_224-233_-specific cells indicating decreased degranulation in these populations (Fig. 6a and b). Interestingly, this defect was notable in PA_224-233_-specific cells but not in NP_366-374_-specific cells. Furthermore, the proportion of cells expressing CD127 was notably higher in PA_224-233_-specific cells compared to NP_366-374_-specific cells in WT mice indicating increased influence of IL-7 on PA_224-233_-specific cells (Fig. 6c).

**Figure 6.**
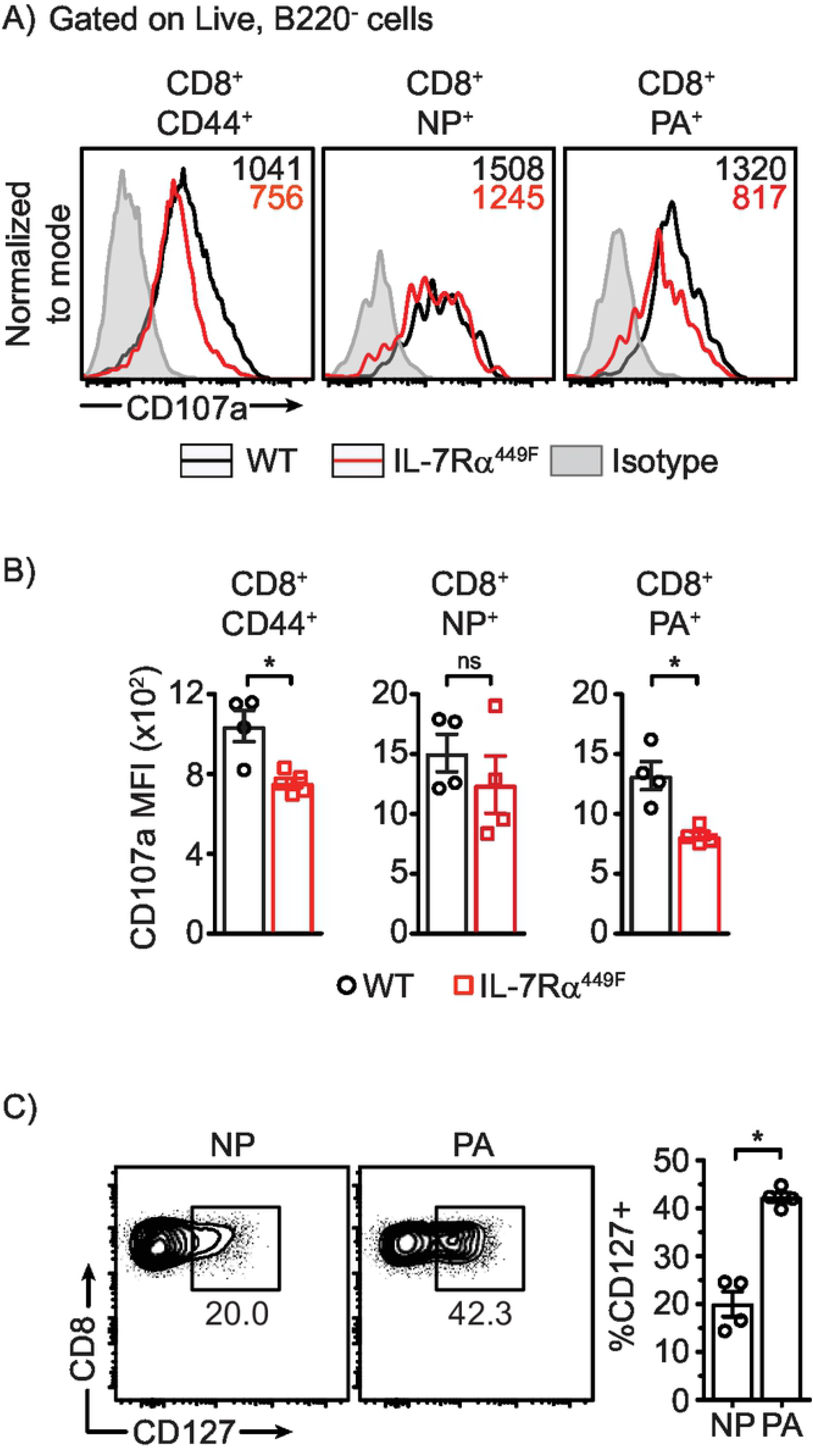
Reduced degranulation of IL-7Rα^449F^ lung CD8 T cells upon re-stimulation. **(a, b)** CD107a expression as median florescence intensity (MFI) in all antigen experienced (CD44^+^), NP_366-374_^+^ and PA_224-233_^+^ CD8 T cells of WT and IL-7Rα^449F^ mice 7 dpi after PMA/Ionomycin re-stimulation. Data presented as **(a)** FACS plots and **(b)** bar graphs. Gated within Live B220^-^, CD8^+^, CD44^+^, NP_366-374_^+^ or PA_224-233_+ cells. Data are representative of two experiments with n=4 per genotype. \**P*<0.05 as determined by two-tailed Student’s t-test. **(c)** Scatter plots and Bar graphs showing flow cytometric analysis comparison of CD127 expression as a percentage within lung NP_366-374_+ and PA_224-233_+ CD8 T cells 7 dpi. Gated within Live B220^-^, CD8^+^, CD44^+^, NP_366-374_^+^ or PA_224- 233_^+^ cells. Data are representative of three experiments with n=4-5 per genotype. \**P*<0.05 as determined by two-tailed Student’s t-test.

To determine if IL-7 signaling affects cytokine production, we treated whole lung single cell suspensions from infected mice with NP_366-374_ and PA_224-233_ peptides *ex vivo* and stained intracellular cytokines to detect IFNγ and TNFα using flow cytometry. NP_366- 374_-specific cells generated low proportion of IFNγ^+^ TNFα^+^ cells compared to PA_224-233_-specific cells regardless of mouse genotype (Fig. 7a). However, WT PA_224-233_-specific cells generated abundant IFNγ^+^ TNFα^+^ cells, which were largely absent within IL-7Rα^449F^ PA_224-233_-specific cells (Fig. 7b). We used TSLPR^-/-^ mice as controls since IL-7Rα is required for both IL-7 and TSLP signaling. We found that TSLPR^-/-^ CD8 T cells presented with reduced cytokine production as well, however, this effect did not follow the same pattern as with IL-7Rα^449F^ mice (Fig. 7a and b).

**Figure 7.**
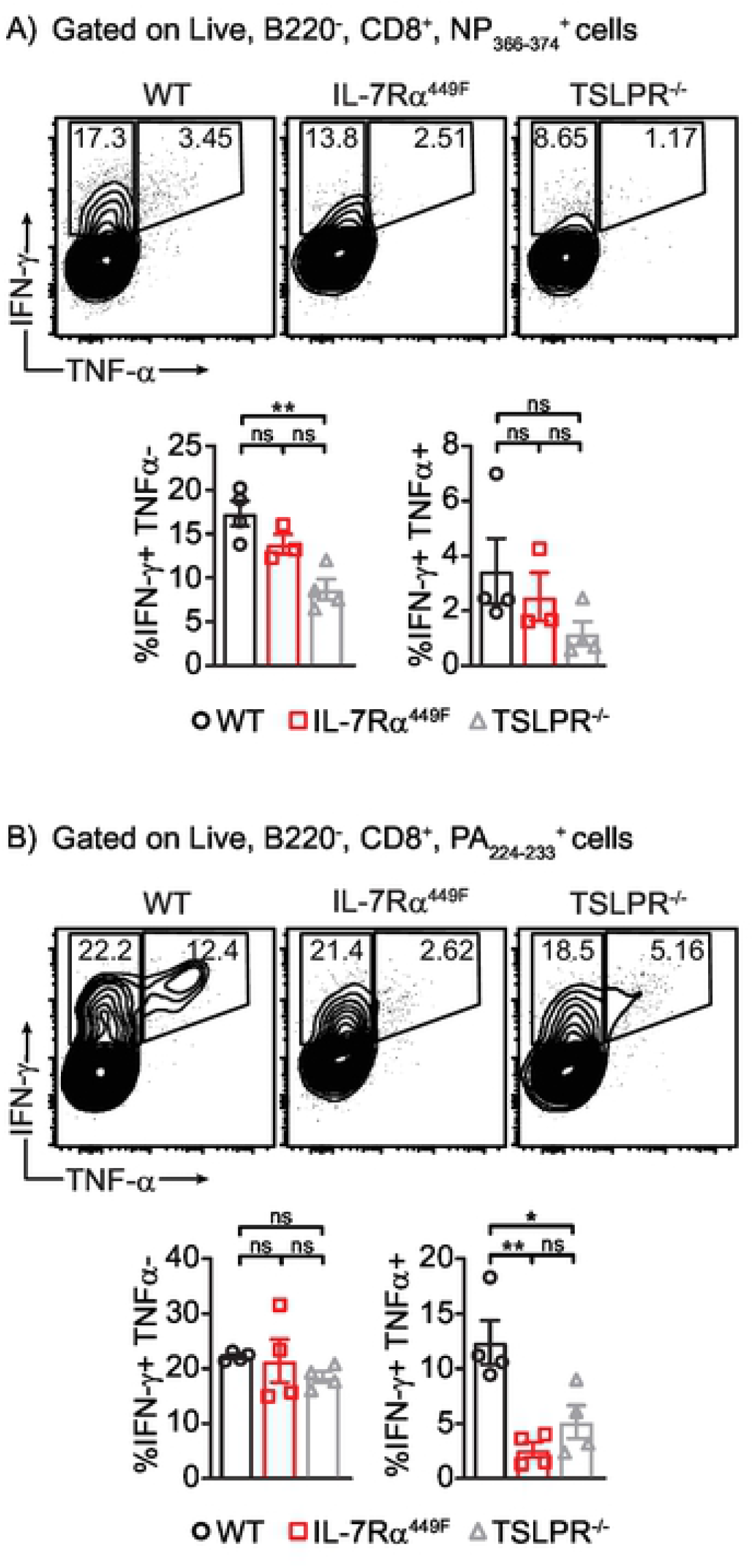
Deregulated cytokine production in IL-7Rα^449F^ and TSLPR^-/-^ lung CD8 T cells. Representative scatter plots and bar charts of IFN-γ^+^ TNF-α^-^ or IFN-γ^+^ TNF-α^+^ CD8 T cells within **(a)** NP_366-374_^+^ and **(b)** PA_224-233_^+^ CD8 T cells 9 dpi and after peptide (NP or PA) re-stimulation. Gated within Live B220^-^, CD8^+^, tetramer^+^ cells. Data is representative of three independent experiments with n=3-5 mice per genotype. \**P*<0.05 as determined by two-tailed Student’s t-test.

To determine if IL-7 independently affects cytokine production, we used IL-7^eGFP/eGFP^ mice that have an eGFP gene inserted disruptively into an *Il7* exon thus serving as an IL-7 ligand knock-out in homozygotes (30). Using this mouse model, we established that IL-7 is required for accumulation of IFNγ^+^ TNFα^+^ cells within PA_224-233_-specific but not NP_366-374_-specific cells (Fig. 8a and b).

**Figure 8.**
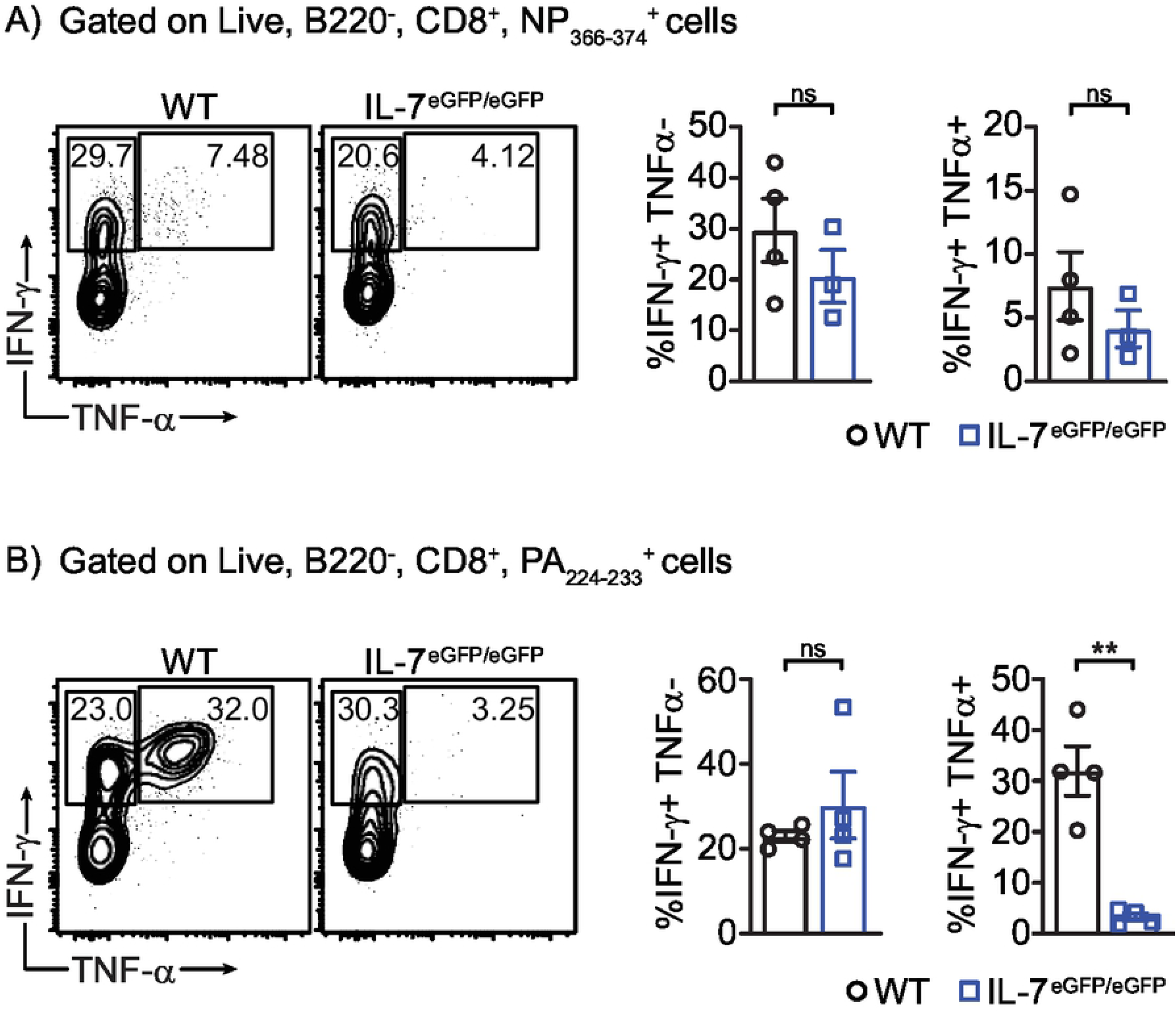
Deregulated cytokine production in IL-7^eGFP/eGFP^ lung CD8 T cells. Representative scatter plots and bar charts of IFN-γ^+^ TNF-α^-^ or IFN-γ^+^ TNF-α^+^ CD8 T cells within **(a)** NP_366-374_^+^ and **(b)** PA_224-233_^+^ CD8 T cells 9 dpi and after peptide (NP or PA) re-stimulation. Gated within Live B220^-^, CD8^+^, tetramer^+^ cells. Data is representative of two independent experiments with n=3-4 mice per genotype. \**P*<0.05 as determined by two-tailed Student’s t-test.

### IL-7 signaling regulates expression of PD-1 in PA_224-233_ but not NP_366-374_-specific CD8 T cells

Expression of inhibitory molecules such as PD-1 is known to be important to negatively regulate T cell activation and limit inflammation by T cells, however, sustained expression of these molecules can lead to dampening of protective immune responses (34). To understand how IL-7 affects PA_224-233_-specific but not NP_366-374_-specific T cells, we evaluated the expression of the inhibitory receptor PD-1. We showed that IL-7Rα^449F^ and IL-7^eGFP/eGFP^ CD8 T cells have higher expression of this molecule (Fig. 9a and b). Specifically, the increase in PD-1 expression in IL-7Rα^449F^ and IL-7^eGFP/eGFP^ CD8 T cells was only evident in PA_224-233_-specific cells but not NP_366-374_-specific cells (Fig. 9a and b). In addition, TSLPR^-/-^ CD8 T cells did not present with increased PD-1 expression (Fig. 9a). Together, this suggests that IL-7 plays distinct roles in CD8 T cell function depending on antigen specificity possibly by regulating PD-1 expression.

**Figure 9.**
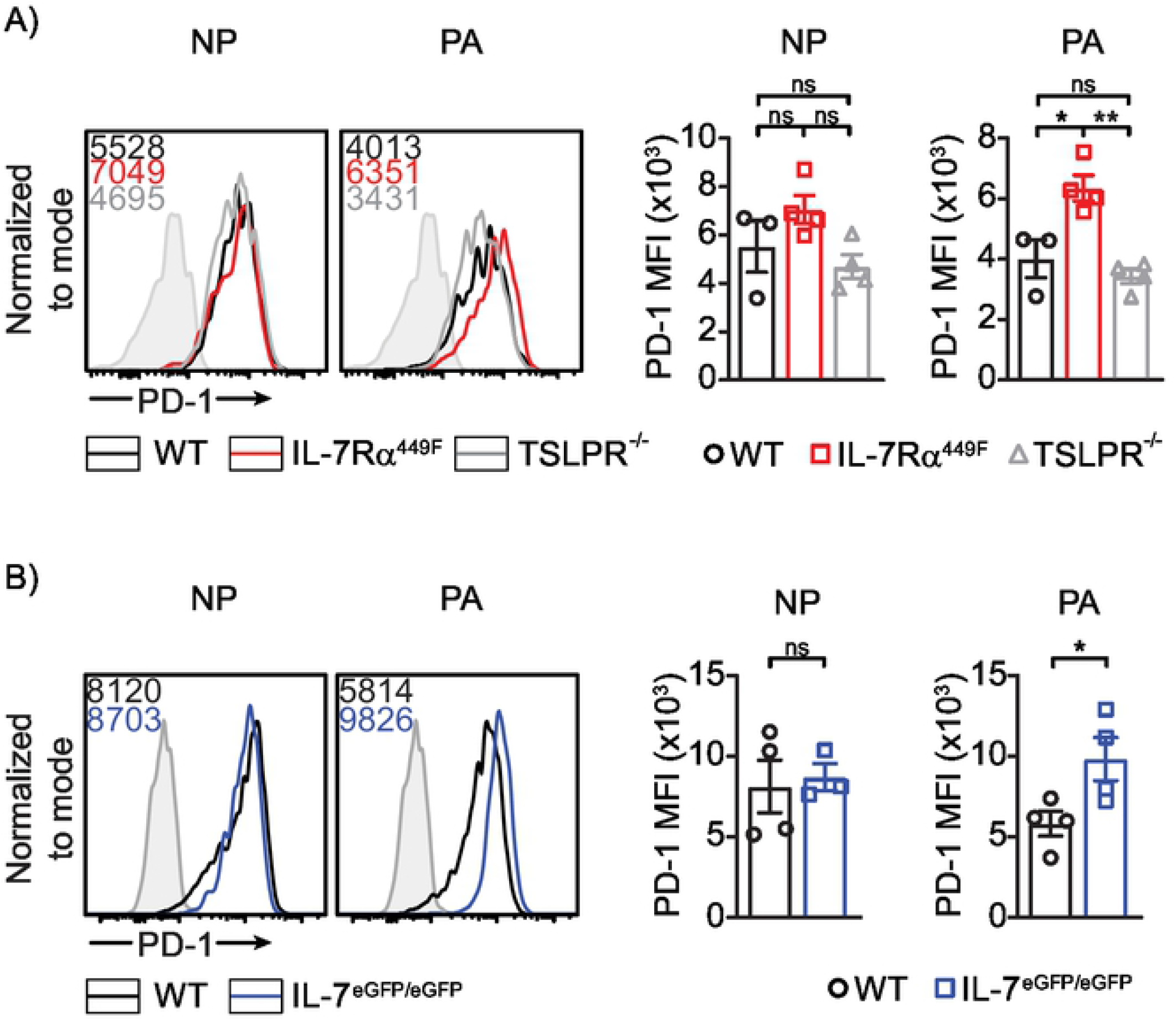
Increased PD-1 expression in IL-7 signaling deficient CD8 T cells. Representative histogram plots and bar charts of PD-1 expression in antigen specific lung CD8 T cells of **(a)** WT vs IL-7Rα^449F^ vs TSLPR^-/-^ and **(b)** WT vs IL-7^eGFP/eGFP^. Gated within Live B220^-^, CD8^+^, tetramer^+^ cells. Data is representative of 2-3 independent experiments with n=3-4 mice per genotype. \**P*<0.05 as determined by two-tailed Student’s t-test.

## Discussion

Initial studies of IL-7 have described its role in B-cell lymphopoiesis and thymopoiesis (35-37). The bone marrow and thymus are the best defined sources of IL-7 production consistent with such roles in primary lymphopoiesis (10). Subsequent studies showed a role for IL-7 in memory cell development and maintenance, in effector response to viral infections and in enhancing T cell functions in chronic conditions (14-16). CD8 T cell expansion and effector function depends on multiple factors including, but not limited to, the cytokine milieu. Previously, we demonstrated that IL-7 is required for the accumulation of tetramer positive CD4 and CD8 T cells during influenza infection (17). The mechanism by which IL-7 accomplishes this and its role in other aspects of T cell response have yet to be elucidated.

We addressed these questions by using mice that express a hypomorphic IL-7Rα (IL-7Rα^449F^) which leads to impaired IL-7 signaling by primarily abrogating STAT5 activation (14). This model provides a better alternative to using IL-7Rα^-/-^ mice since IL-7Rα^449F^ mice have defective signaling yet retain sufficient number of T cells to perform infection studies. We have previously used this mouse model to demonstrate an intact CD8 T cell effector response to intracellular *Listeria monocytogenes* infection (14). In the current study, we found that defective IL-7Rα signaling led to reduced accumulation of influenza-specific CD8 T cells in the secondary lymphoid organ (mdLN) at early priming stages (5 dpi) which ultimately led to reduced accumulation of influenza-specific CD8 T cells in the lungs. Examination of IL-7Rα^449F^ mdLN revealed a great reduction in its size. This is consistent with the fact that IL-7 is required for the development of lymphoid tissue inducer (LTi) cells that seed LN anlagens and drive the organogenesis of LNs (19). Considering IL-7Rα^449F^ mice had reduced, albeit notable, numbers of influenza-specific CD8 T cells in their lungs, it is unclear where and how these CD8 T cells expand to significant numbers with an abnormal mdLN. It is possible that tertiary lymphoid organs in the lung tissues such as inducible bronchus-associated lymphoid tissue (iBALT) provide a suitable environment for the accumulation of *de novo* pathogen specific cells without requiring IL-7 or LTi cells (38). We have shown that despite such extrinsic factors, IL-7Rα signaling is required cell intrinsically by CD8 T cells for early priming in the mdLN.

It is known that a population of CD8 T cells specific to a distinct peptide do not originate from a single naïve precursor but rather from 10s to 100s of precursors (39-41). IL-7 signaling deficient mice have reduced thymic output of T cells, and this may result in a more stochastic or reduced chance of a T cell encountering a cognate MHC-peptide leading to reduced clonal expansion. In addition to these effects, IL-7 can play a role cell intrinsically by affecting TCR repertoire via VDJ recombination or TCR sensitivity (8, 42). We addressed this by adoptive transfer of CD8 T cells bearing a transgenic TCR (OT-I) in equal numbers (WT vs IL-7Rα^449F^) intravenously into congenic WT mice. Using this approach, we found that OT-I;IL-7Rα^449F^ CD8 T cells expanded in response to infection with influenza PR8-OVA to a lower extent compared to OT-I CD8 T cells within total host CD8 T cells. Our findings show that IL-7 is intrinsically important for the accumulation of pathogen-specific CD8 T cells during early priming phase in the mdLN independent of TCR specificity and the number of naïve T cell precursors. We have previously shown that IL-7Rα^449F^ CD8 T cell form antigen-specific cells normally during systemic *in vivo* infection with *L. monocytogenes* yet do not proliferate well when exposed to suboptimal TCR stimulation *in vitro* in contrast to high dose TCR stimulation (14). It is possible a low dose, local influenza infection recapitulates the low level TCR stimulation model whereby IL-7 plays an essential role in CD8 T cells under low TCR avidity activation.

In addition to the intrinsic role that IL-7 plays in CD8 T cells, we found that IL-7Rα^449F^ mice have continued accumulation of CD11b^+^ DCs in the lungs while in WT mice, the number of CD11b^+^ DCs peaks at 7 dpi then subsides. Previous studies using IL-7^-/-^ and IL-7Rα^-/-^ showed normal development of DC precursors in the BM, however these mice had reduced migratory DCs in secondary lymphoid organs (22). Our BM chimera experiments showed that the effect of IL-7 in DC accumulation was indirect. Therefore, the accumulation of DCs we noted in IL-7Rα^449F^ mice was not due to a problem with migration or maturation and was likely due to the fact that viral clearance was impaired which led to continued recruitment of DCs to the lungs.

We have demonstrated for the first time that IL-7 is inducible in lung epithelial cells in response to viral infection *in vivo*. While the increase in total IL-7-eGFP^+^ cells in the lung was modest, we noted a shift in the population that are positive for IL-7-eGFP. In naïve mice, the majority of IL-7-eGFP^+^ cells were lymphatic endothelial cells (LECs) as previously reported (30, 31). However, following infection with influenza, IL-7-eGFP^+^ epithelial cells (ECs) and to a lesser extent stromal cells (SCs) expanded while LECs decreased in frequency. Our results demonstrate that IL-7 can be produced by lung tissues and this could shape the function of CD127 expressing CD8 T cells locally. It is unclear to what extent IL-7 produced by epithelial cells influences nearby cells and the significance of the shift in sources of IL-7. In addition, more sensitive approaches are required to compare the levels of IL-7 expression between the different lung tissues.

Terminal differentiation of activated CD8 T cells is important for the generation of short-lived effector cells (SLECs) that express killer cell lectin-like receptor G1 (KLRG1) and low CD127 (6, 32). We found that following infection with influenza, IL-7Rα^449F^ pathogen-specific CD8 T cells have reduced expression of KLRG1 and terminal differentiation to SLEC. The difference in KLRG1 expression between WT and IL-7Rα^449F^ mice was greater within PA_224-233_-specific cells than in NP_366-374_-specific cells. We also noted a similar trend in CD107a expression between WT and IL-7Rα^449F^ mice whereby the defect in CD107a expression was more pronounced in PA_224-233_-specific cells than in NP_366-374_-specific cells. Increased expression of CD127 by PA_224-233_-specific cells supports the hypothesis that PA_224-233_-specific cells have elevated dependence on IL-7 signaling.

Pro-inflammatory cytokines such as IFNγ and TNFα are important during an anti-viral response to help with recruiting and activating other cells. Within NP_366-374_-specific cells, we did not observe any differences in expression of IFNγ and TNFα between WT and IL-7Rα^449F^ or IL-7^eGFP/eGFP^ CD8 T cells. However, the large reduction in IFNγ^+^ TNFα^+^ population in PA_224-233_-specific cells of IL-7Rα^449F^ or IL-7^eGFP/eGFP^ mice followed a similar pattern to our findings with CD107a. NP_366-374_-specific cells did not generate IFNγ^+^ TNFα^+^ cells as notably as PA_224-233_-specific cells, consistent with other groups (43, 44). Our work further corroborates what previous studies have shown, that TSLP shapes effector T cell responses following influenza infection. However, these effects were shown to occur indirectly through programming of DCs (45). Altogether, this suggests that IL-7 may differentially regulate effector function of pathogen-specific CD8 T cells between T cell clones. We have not determined the molecular mechanism and downstream signaling to understand how this occurs.

Previous studies have shown hierarchical differences between NP_366-374_ and PA_224-233_-specific CD8 T cells whereby tissue resident memory CD8 T cells that are NP_366-374_-specific expressed higher levels of inhibitory molecules including PD-1 at 30 dpi and beyond due to persistent antigen exposure and TCR stimulation (46). IL-7 is also known to enhance cytokine production and reverse T cell exhaustion by repressing inhibitory pathways during chronic viral infections in mice (15, 16). It is well established that TCR signaling duration correlates positively with PD-1 expression but little is known about this phenomenon in the context of influenza-specific CD8 T cells (47, 48). In an acute hepatitis B virus infection, PD-1 expression in CD8 T cells is negatively correlated to CD127 expression, and blocking PD-1 in acute lymphocytic choriomeningitis virus infection increases the frequency of the CD127^+^ population (49, 50). Our finding that PD-1 expression is higher within PA_224-233_-specific but not NP_366-374_-specific CD8 T cells with IL-7 signaling deficiency is indicative of an antigen-dependent role for IL-7 in regulating their function. This is suggestive of a negative regulatory role for IL-7 on PD-1 expression dependent on TCR clonotype. This hypothesis is corroborated by a previous study where infection with a high pathogenicity influenza/A virus strain, such as PR8, resulted in elevated PD-1 expression in antigen-specific CD8 T cells compared to infection with the low pathogenicity influenza/A x31 strain (44). This in turn inhibited effector function by specifically affecting development of IFNγ^+^ TNFα^+^ cells (44). Our studies suggest that PA_224-233_-specific IL-7 signaling deficient CD8 T cells do not receive the necessary signals to down regulate PD-1. Further studies are required to define the relationship between IL-7 and PD-1 in an acute infection setting and the mechanism by which this specifically affects T cells in a clone-specific manner.

In summary, we have found that IL-7 is required for an optimal response to acute influenza infection as it shapes the early priming stages of CD8 T cells. Moreover, IL-7 produced by lung tissues is important for the terminal differentiation and effector function of CD8 T cells in specific TCR clones of CD8 T cells. Various cytokines have the ability to enhance CD8 T cell responses, however, rigorous testing is necessary to evaluate the adverse responses that these cytokines have on bystander cells. Using cytokines such as IL-7 to complement existing therapies may be beneficial given fewer off target effects due to the limited subset of cells that express CD127. IL-7 is currently in clinical trials for treatment of infections and tumors. Additional studies are necessary to expand the use of IL-7 in other conditions and to study its efficacy when delivered in combination with other agents.

## Methods

### Mice

All mice were housed and used in the Center for Disease Modelling facility (CDM) at the University of British Columbia (UBC) and all work with animals was carried out with approval and in accordance with the ethical guidelines of the University of British Columbia Animal Care and Biosafety Committees. IL-7Rα^449F^ mice were generated in-house as described (14). Briefly, they express a mutant form of the IL-7Rα with a single amino acid mutation from Tyr to Phe at position 449. C57BL/6, BoyJ (B6.SJL-Ptprca Pepcb/BoyJ) and C57BL/6-Tg (TcraTcrb) 1100Mjb/J (OT-I) mice were obtained from the Jackson Laboratory (Bar Harbour, ME, USA). IL-7^eGFP/eGFP^ mice were a gift from J.M. McCune (UCSF) (30). In all cases, age-matched and sex-matched male and female mice between the ages of 6-12 weeks were used.

### Virus

Influenza A/PR/8/34 (PR8) was purchased from Charles River Laboratories (Wilmington, MA). Influenza A/PR/8/34-OVA (PR8-OVA) was propagated in-house in chicken eggs as previously described (51). Mice were sub-lethally infected under anesthesia (isoflurane) with 5 Hemagglutinin Units (HAU) of influenza PR8 or 64 HAU of influenza PR8-OVA in 12.5 µL of sterile PBS intranasally.

### Tissue preparation

Mice were anesthetized with 5% isoflurane in 1L/min O_2_ and euthanized by cervical dislocation and perfused with 10 ml cold PBS (+5%FBS, 2mM EDTA). Lungs were excised and processed by mincing with scissors followed by enzymatic digestion using 180 units/ml collagenase IV and 20 µg/ml DNase I (Worthington biochemical LS004188 and LS002139) in 5 ml RPMI incubated at 37°C for 30-45 mins in a shaker incubator before filtering through 70 µm filters and lysing RBCs with ACK lysis buffer. To assess non-hematopoietic cells in the lungs, dispase (1u/ml) was added to the enzyme cocktail. Mediastinal lymph nodes (mdLN) were collected and crushed through 70 µm filters and suspended as single cells in cold PBS (5%FBS, 2mM EDTA).

### Antibodies and Flow cytometry

All cell surface staining was done at 4°C for 30 mins in the dark. Anti-CD8a [53-6.7] (APC-eFluor780), anti-B220 [RA3-6B2] (PE-eFluor610), anti-CD44 [IM7] (PE-Cy7), anti-IFNγ [XMG1.2] (A488), anti-TNFα [MP6-XT22] (PE), anti-MHCII [M5/114.15.2] (FITC), anti-CD11b [M1/70] (PE-Cy7), anti-CD326/EpCAM [G8.8] (PE-Cy7) anti-CD107a [eBio1D4B] (PE) and Rat IgG2a kappa Isotype control [eBR2a] (PE) were purchased from Thermo Fisher (Waltham, Massachusetts). Anti-KLRG1 [MAFA] (APC), anti-CD127 [SB/199] (PE) anti-PD-1 [29F.1A12] (BV510), anti-CD11b [M1/70] (PE-Cy7), anti-CD11c [N418] (biotin), anti-CD45 [30-F11] (Pacific Blue), anti-CD45.2 [104] (PerCP/Cy5.5 or BV421), anti-CD31 [MEC13.3] (biotin), anti-Gp38 [8.1.1] (PE) and anti-F4/80 [BM8] (PE-Cy7) were purchased from Biolegend (San Diego, California). Anti-CD103 [M290] (PE), anti-TCR Vα2 [B20.1] (PE) and anti-CD45.1 [A20] (APC) were purchased from BD Biosciences (Franklin Lakes, New Jersey). Anti-CD11c [N418] (Alexa fluor-647), anti-CD45.1 [A20] (A488), anti-B220 [RA-6B2] (FITC) and anti-F4/80 [BM8] (biotin) were purchased from AbLab (Vancouver, British Columbia).

Tetramer staining was done at room temperature for 30 mins in the dark. H2-K^b^ tetramers loaded with immune-dominant NP_366–374_ and PA_224–233_ peptides from influenza and labeled with Brilliant Violet-421 or Alexa fluor-647 were manufactured and donated by the NIH Tetramer Core Facility (Atlanta, GA).

Viability staining [cat# L34957 and 65-0865-14] (Thermo Fisher) was used according to manufacturer’s instructions.

Samples were collected on either a FACSCanto, LSRII (BD Biosciences) or the Attune NxT (Thermo Fisher) and data were analyzed with FlowJo software Tree Star (Ashland, Oregon).

### Bone marrow chimeras

Recipient mice were irradiated with 2 doses of 6.5 grey (Gy) or 650 rad at least 4 hours apart. For the following 10 days, they were supplemented with antibiotics ad libitum (2mg/ml neomycin sulfate). 24 hours after radiation, femurs and tibias were collected from donor CD45.1 and CD45.2 mice (WT and IL-7Rα^449F^ respectively). RBCs were removed using sterile ACK lysis. For tetramer response experiments, a total of 1×10^6^ donor bone marrow (BM) cells were injected intravenously (I.V.) at a 1:1 ratio to deliver WT:WT or 1:10 ratio to deliver WT:IL-7Rα^449F^ into Rag1^-/-^ hosts. For dendritic cell experiments, a total of 5×10^5^ donor BM cells were injected I.V. at a 1:1 ratio to deliver WT:WT or WT: IL-7Rα^449F^ into C57BL/6J;Boy/J (CD45.1/.2) hosts. 6-8 weeks elapsed for reconstitution before challenge with influenza infection. After euthanasia, spleens and BMs were assessed for reconstitution efficiency and ratios.

### Adoptive transfer

Single cell suspensions were prepared from multiple OT-I and OT-I;IL-7Rα^449F^ mice spleens and CD8 T cells were purified using the CD8 T cell negative selection kit (EasySep™ Mouse CD8+ T Cell Isolation Kit) from Stem Cell Technologies. 1×10^6^ cells were transferred I.V. into BoyJ (CD45.1) hosts and 24 hours later hosts are challenged with 64HAU PR8-OVA intranasally. MdLN was harvested at experimental endpoint and cells were stained for surface markers and analyzed by flow cytometry as described above.

### Cell culture and *II-7* RT-qPCR

A549 (ATCC CCL-185) human type II alveolar epithelial cells were obtained from the American Type Culture Collection (ATCC) (Manassas, Virginia). Cells were passaged and expanded in 10% FBS F-12K Medium from ATCC (Cat No. 30-2004). For experimental use, 5×10^5^ A549 cells were seeded into 6-well plates in media and expanded for 24 hours to achieve confluence. After 1 hour of serum starvation, cells were infected with 200 HAU PR8 in PBS and incubated for 1 hour on a plate shaker to initiate infection. Virus containing PBS was aspirated and replaced with F-12K media containing 0.5% BSA and 0.5 µg/mL N-tosyl-L-phenylalanine chloromethyl ketone (TPCK) treated trypsin. Cells were then incubated for the assigned experimental time points. Cells were lysed and RNA was extracted using PureLink RNA Mini Kit (Thermo Fisher). After treatment with amplification grade DNase I (Thermo Fisher), cDNA was generated using the iScript cDNA synthesis kit (Bio-Rad) and cDNA quantification was performed using the Ssofast EvaGreen Supermix kit (Bio-Rad). Primer sequences are as follows. β-actin Forward: GAC ATG GAG AAA ATC TG; β-actin Reverse: ATG ATC TGG GTC ATC TTC TC; Human IL-7 Forward: CCA GGT TAA AGG AAG AAA ACC; Human IL-7 Reverse: TTT CAG TGT TCT TTA GTG CC; Human IFN-β Forward: ACG CCG CAT TGA CCA TCT AT; Human IFN-β Reverse: GTC TCA TTC CAG CCA GTG CTA; M1 Forward: AGA TGA GTC TTC TAA CCG AGG TCG; M1 Reverse: TGC AAA AAC ATC TTC AAG TCT CTG. Measurements were acquired using the CFX96 Touch Real-Time PCR Detection System (Bio-Rad).

### *Ex vivo* T cell re-stimulation

Lungs from mice infected (7-9 days) with PR8 were excised and prepared as above. To measure CD107a, 5×10^6^ lung cells were re-stimulated for 4 hours (37°C, 5% CO_2_) with 1% BSA RPMI containing 50 ng/ml PMA and 500 ng/ml Ionomycin from Sigma-Aldrich (St. Louis, Missouri); Monensin from BD Biosciences (Franklin Lakes, New Jersey) used according to manufacturer’s instructions; and anti-CD107a [eBio1D4B] PE (33). Following re-stimulation, cells were stained for viability and surface markers then analyzed by flow cytometry.

To measure IFN-γ and TNF-α, 5×10^6^ lung cells were re-stimulated for 3 hours (37°C, 5% CO_2_) with 10 nM NP_366–374_ and PA_224-233_ peptides from Anaspec (Fremont, California) in the presence of Brefeldin/A from BD Biosciences (Franklin Lakes, New Jersey) in 1% BSA RPMI. Following re-stimulation, cells were stained for viability and surface markers followed by intracellular cytokine staining using the Cytofix/Cytoperm kit from BD Biosciences (Franklin Lakes, New Jersey) then analyzed by flow cytometry.

## Author contributions

AS conceived and designed the project, performed and analyzed experiments, and wrote the manuscript. JJ, HBS and CY performed and analyzed experiments and reviewed the manuscript. JS performed and analyzed experiments. NA conceived and designed the project and reviewed the manuscript.

## Acknowledgements

We would like to thank Adam Plumb, Julia Lu and Etienne Melese for critical revision of this manuscript. We would also like to thank the staff at the Centre for Disease Modeling for animal husbandry and the UBC Flow Cytometry Facility.

## Figure legends

Supplementary Figure 1. Expansion of adoptively transferred OTI-IL-7Rα^449F^ CD8 T cells is impaired in the mdLN following influenza infection as early as 3 dpi. Scatter plot and representative bar graph of CD45.2^+^ Vα2^+^ CD8 T cells. Gated within Live B220^-^ CD8^+^ cells. Datum is representative of a single experiment with n=4. \*\**P*<0.01 as determined by two-tailed Student’s t-test.

Supplementary Figure 2. Loss of IL-7Rα signaling leads to increased accumulation of CD11b^+^ CD103^-^ dendritic cells in the lungs. **(a)** Flow cytometric analysis showing total number of CD11b^+^ CD103^-^ (left) and CD11b^-^ CD103^+^ (right) dendritic cells in the lungs of WT and IL-7Rα^449F^ mice at indicated days post infection presented as a bar graph. Gated within Live CD45^+^, B220^-^, F4/80^-^, CD11c^hi^, MHCII^hi^, CD11b^+/-^ and CD103^-/+^. **(b, c)** Bone marrow chimera analysis of lung CD11b^+^ CD103^-^ and CD11b^-^ CD103^+^ dendritic cells presented as **(b)** bar graphs and **(c)** FACS plots. **(b)** Data presented as ratio of the CD45.2:CD45.1 and plotted with log_10_ transformation to normalize skewed data points. Data are representative of two experiments with n=4-6 per genotype. \*\*\**P*<0.001 as determined by two-tailed Student’s t-test.

